# Evolution on degenerate fitness landscapes is not neutral: curvature drives directional bias

**DOI:** 10.64898/2026.02.14.705886

**Authors:** Razi Fachareldeen, Naama Brenner

## Abstract

Degeneracy - the multiplicity of phenotypes with equivalent fitness - is a prevalent feature of biological systems. Such degeneracy is often associated with neutral evolution, under the assumption that adaptive dynamics on degenerate fitness manifolds lacks direction. Here we show that this is not generally the case. Using a minimal model of evolutionary dynamics on smooth degenerate fitness landscapes, we demonstrate that stochastic mutation-selection dynamics induce a directional drift on manifolds of optimal fitness toward regions of reduced curvature. This drift arises from an interaction between population variability and landscape curvature: curvature shapes phenotypic variation, which in turn biases evolutionary exploration even when fitness gradients vanish. As a result evolution exhibits an *implicit bias*, preferentially selecting flat and robust regions of the degenerate fitness manifolds, without explicit optimization for these properties. Interestingly, similar flatness-seeking implicit biases have been discovered in other stochastic optimization algorithms; here we reveal their different underlying mechanisms despite similar outcome, highlighting the unique properties of evolutionary dynamics. Our results highlight a general mechanism by which degeneracy shapes long-term evolutionary outcomes, affecting our interpretation of phenotypic variability, robustness, and neutrality in high-dimensional biological systems.

## A. Introduction

Understanding how evolution operates on complex fitness landscapes remains a central challenge in evolutionary biology [1]. Classical intuition often portrays adaptation as hill climbing toward isolated fitness peaks, with fitness gradients providing clear evolutionary direction [2]. However, accumulating empirical and theoretical evidence indicates that real high-dimensional biological landscapes are highly degenerate. For example, studies at the genome sequence level in *E. coli* [3] and *C. servisea* [4, 5], analyses of diverse Genotype-Phenotype maps [6], and models of cell metabolism [7] have consistently revealed that biological landscapes are smoother and more navigable than expected, and contain large degenerate manifolds of optimal or near-optimal fitness.

Degenerate fitness landscapes are commonly associated with neutral evolution, in analogy with random changes in the frequencies of alleles with equal fitness [8]. When fitness gradients vanish along extended manifolds of equally fit phenotypes, evolutionary dynamics may be expected to proceed as undirected diffusion, shaped primarily by historical contingency and stochastic fluctuations (Fig. 1(a)). Here we challenge this view and show that evolution on degenerate smooth fitness landscapes is generically *not* neutral. Instead, mutation-selection dynamics induce directional biased motion along manifolds of optimal fitness, even in the absence of fitness gradients. This motion arises from an interaction between population-level variability and local curvature of the fitness landscape: curvature suppresses variation in sharp directions while allowing it to persist in flatter ones. Over evolutionary time, this bias produces an effective drift toward regions of reduced curvature, where phenotypes are intrinsically more robust to perturbations (Fig. 1(b)).

**FIG. 1.**
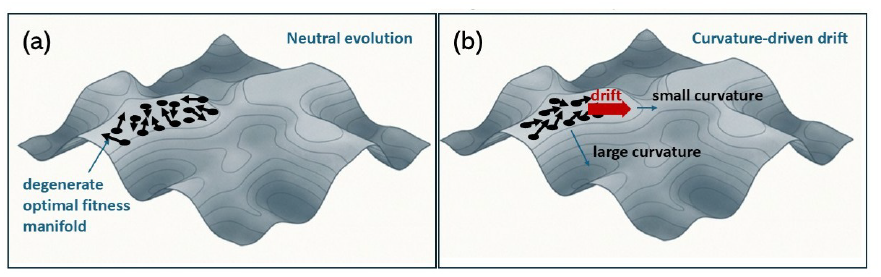
Evolution on degenerate landscapes is not neutral. A complex fitness landscape features a degenerate manifold of maximal fitness (lightest shade). In the absence of gradients, a naive expectation would be that evolution is neutral on this manifold **(a)**, and that individuals who have reached it would move stochastically. However, we show that this is not the case if curvature is not constant across the manifold **(b)**; a deterministic drift emerges from the interplay of changing curvature and population variability, towards the flatter regions on the manifold.

To illustrate this mechanism, we analyze evolutionary dynamics on a minimal, analytically tractable fitness landscape that captures two essential features of realistic biological systems: extended degeneracy and heterogeneous curvature. We show that populations rapidly approach the manifold of optimal fitness and subsequently drift along it toward flatter regions. This behavior does not rely on explicit selection for robustness, nor on fitness differences along the manifold, but instead emerges from second moment statistical effects intrinsic to mutation-selection dynamics.

Closely related phenomena have recently attracted significant attention in machine learning, where optimization of neural networks reveals extended manifolds of near-degenerate solutions in loss landscapes. Empirically, stochastic gradient descent (SGD) - the workhorse algorithm of modern machine learning - exhibits an implicit bias toward flat minima, a property associated with robustness and generalization [9–13]. Utilizing a physical model of SGD, we compare the dynamics of an ensemble of optimizers with that of an evolving population. We find that the interplay of variability with landscape is very different in the two types of dynamics; nevertheless both exhibit a tendency to drift toward flatter regions of degenerate manifolds. This highlights the generality of degeneracy as a regime where curvature can interact with stochasticity in different ways to shape long-term outcomes across distinct adaptive systems.

Our results have broad implications for how evolved biological systems are interpreted. They suggest that phenotypic robustness, which characterizes flat regions of the fitness landscape, can arise as an emergent property of long-term evolutionary dynamics without requiring its direct selection. Likewise, phenotypic variability observed in natural populations need not reflect noise or lack of optimization, but may instead indicate residence within flat regions of the landscape, where states are similarly fit.

## B. Directional bias on general fitness landscapes

Does mutation-selection dynamics on degenerate manifolds of equal fitness necessarily lead to neutral diffusion? To address this question, we first consider evolutionary dynamics at a general level, without specifying a particular fitness landscape. We show that when curvature varies, evolution undergoes a directional bias towards flatter regions, in addition to gradient climbing. We use a minimal model of evolutionary dynamics and derive a general one-generation mapping. The population is described by *p*(x, *t*), a probability density in a continuous state **x** at time *t*. It proceeds in discrete non-overlapping generations with two operators acting on the distribution: the mutation operator 𝕄 spreads the population by adding an independent noise to each state; the distribution undergoes a convolution with a Gaussian kernel 𝒩 (0, **Σ**_𝕄_)

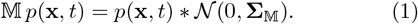

The selection operator S reshapes the distribution based on relative fitness:

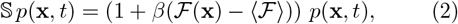

where *β* is the selection strength and ⟨ℱ ⟩ the average population fitness (alternative formulations of the selection operator are discussed in the Supplementary Material, Section 1.2). Over one generation,

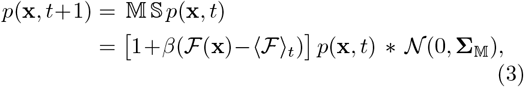

where ⟨·⟩_*t*_ is an average over the distribution *p*(**x**, *t*).

To leading order in population variability (or equivalently, assuming the distribution is Gaussian), the average state of the population advances over one generation as

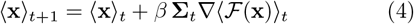

It proceeds along the *natural gradient* [14] of the smoothed fitness landscape. The natural gradient is a potentially anisotropic rescaling of the gradient by the population covariance matrix **Σ**_*t*_. As evolution can only advance where there is variance to act upon, this rescaling reflects the evolvability potential in different directions of state space.

To see the effect of averaging the landscape over the population distribution, we perform the average explicitly and find that it induces a bias that effectively adds an extra term to the fitness function being optimized:

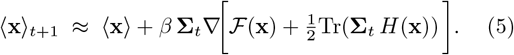

The term Tr(**Σ**_*t*_*H*) couples the population variability with local landscape curvature, and adds an effective term - an implicit bias - to the function under the gradient. It is a variance-weighted average of curvature, negative near fitness maxima and more negative in regions of high curvature; therefore in terms of optimization, it places a penalty on such regions. Differences in this penalty induce a driving force for the motion of the population average towards flatter regions, in addition to the fitness gradient.

Detailed derivations and extensions of the results in this section are given in Supplementary Material Section1. Next, we illustrate this general phenomenon on an analytically tractable fitness landscape, where the consequences of curvature-variance coupling can be computed explicitly.

## C. Directional bias on a toy model landscape

For realistic biological landscapes in high dimensions, degenerate manifolds are abundant, gradients can vanish and the regularization effect described above becomes a major force for evolution. However, such landscapes are difficult to describe or visualize. A similar property also appears in the loss landscapes of artificial neural networks; useful models of such landscape have been developed in the Machine Learning (ML) literature [11, 13, 15]. Following this line of work, we expand the fitness function ℱ in *d* dimensions around the maximal ridge to second order:

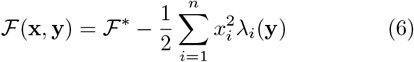

where ℱ^∗^ is the optimal value, the vector **x** ∈ ℝ^*n*^ represents coordinates in the non-degenerate subspace (orthogonal to the optimal ridge), and **y** ∈ ℝ^*d*−*n*^ - the coordinates in the degenerate subspace (along the optimal ridge). The curvatures *λ*_*i*_ generally vary with position **y** along the degenerate manifold, implying that although all points are degenerate in terms of their fitness, their curvature varies. We will focus on the simple two-dimensional case

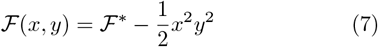

where the maximal fitness ℱ^∗^ is obtained on the manifold *xy* = 0. Despite its simplicity, this two-dimensional landscape maintains the key features described above: a continuous manifold of degenerate connected maxima, with a varying curvature in orthogonal directions. Specifically, the manifold becomes progressively flatter as one approaches the origin (0, 0), which is the point of minimal curvature.

Simulations of the evolutionary dynamics Eq. (3) on this landscape were performed with isotropic mutation kernel *σ*^2^**I**. Fig. 2(a) shows the positive quadrant of the (*x, y*) plane with gray contour-lines of the landscape, and population average trajectories with different mutation strengths (different colored lines). For large *σ*, the smoothing effectively lifts the degeneracy from the optimal manifold, and the flattest point (0, 0) becomes the global minimum; the average trajectory then follows the gradient directly to that point (brown line). For small *σ* there is still near-degeneracy on the optimal line *x* = 0, and the trajectories can be decomposed into two parts: first, achieving optimal fitness on the manifold; second, upon reaching the optimal manifold, the average proceeds to drift along the manifold, directed towards the flattest point (0, 0) (light orange lines). Fig. 2(b) shows the curvature at the population average, decreasing along time, with the same colors corresponding to different mutation strengths.

**FIG. 2.**
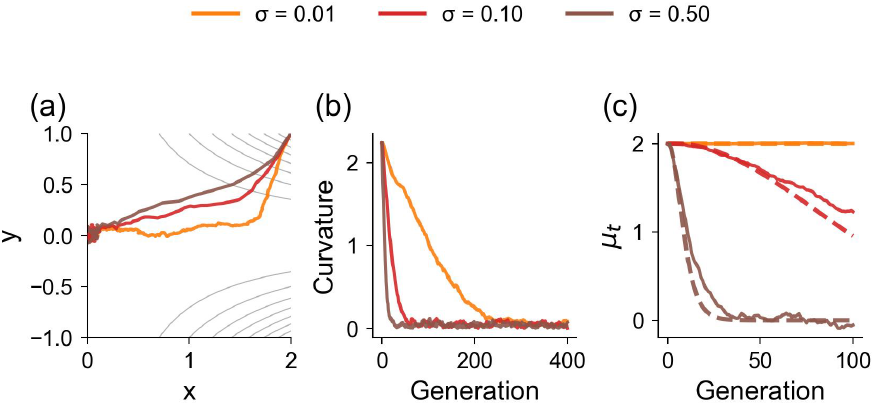
Evolutionary dynamics on a two-dimensional fitness landscape with a degenerate optimal manifold. The mutation kernel is Gaussian with covariance **Σ**_𝕄_ = *σ*^2^**I**. Population averages under mutation-selection dynamics, for three values of increasing mutation strength. **(b)** Curvature Tr(**H**( ⟨**x⟩** _*t*_)) evaluated at the population mean along the same trajectories. **(c)** A population starts with an average on the optimal degenerate manifold, ⟨x⟩ _0_ = (*µ*_0_, 0)^⊤^ = (2, 0). Remaining on *y* = 0, the *x*-coordinate of its average exhibits a drift towards the flattest point on the manifold (simulation - solid lines), as predicted by theory (Eq. 4 - dashed lines). In all panels the election coefficient *β* = 1; the initial covariance matrix **Σ**_0_ = *σ*^2^**I**; and population size 1,000. Panels (a) and show 400 generations; panel (c) shows 100 generations.

The drift can be calculated exactly by starting the population at an initial condition on the optimal manifold, with an average ⟨**x**⟩ _*t*_ = (*µ*_*t*_, 0)^⊤^ and a diagonal covariance matrix with elements *a*_*t*_, *b*_*t*_, denoted as **Σ**_*t*_ = diag(*a*_*t*_, *b*_*t*_). The population average proceeds along the *x*-axis as

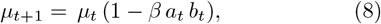

while the average in the *y* direction remains zero. The drift velocity increases with *µ*_*t*_, the *x*-value of the average, where the curvature is larger; it increases with higher population variances, and with stronger selection. Fig. 2(c) shows how this analytic result compares with simulated dynamics. For more details see Supplementary Material Section 2.2.

## D. Curvature shapes phenotypic variability and induces drift

What is the dynamic mechanism underlying the directed bias on the flat manifold? To answer this question we examine the evolution of the population covariance matrix. Under the Gaussian approximation, we can also derive the one-generation update rule for this covariance (see Supplementary Material, Section 1). To leading order in *β*,

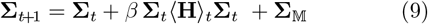

were ⟨H⟩ _*t*_ is the Hessian of the fitness landscape av
eraged over the population distribution. In regions near fitness maximum where Hessian is negative, the variance along all directions is reduced, and more so along steep directions, shaping the covariance to extend relatively more in flatter directions. In regions near a fitness minimum, where curvatures are positive, the covariance is extended, and more so towards the steeply increasing direction. This interplay induces an effectively landscape-dependent variability; therefore, although the **noise** injected into the system by mutations may be isotropic (or more generally, blind to its fitness outcome), the integrated outcome of mutation and selection over the history of the population renders the **variability**, which is the substrate for selection, fitness-dependent.

The last term in the update is the additive mutation covariance. In the approximation where mutations act as a convolution, the covariance is additive,

To see how the covariance evolves in time more concretely, we return to our two-dimensional landscape and start the population on the degenerate optimal manifold as before. Here we take a diagonal mutation covariance matrix with elements **Σ**_𝕄_ = diag(*m*_*x*_, *m*_*y*_). The average Hessian takes the form 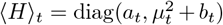 the population covariance commutes with the Hessian, and thus continues to evolve as a diagonal matrix with elements

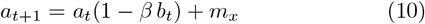

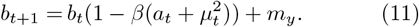

Starting from an initial condition far from the origin, where 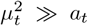, the variance in the sharp *y*-direction rapidly decreases, distorting the population to form an anisotropic distribution, still aligned with the Hessian but with variance much larger along the flat *x*-direction. This distortion is driven by the large gap between the curvatures in the two directions and propels the average drift. A comparison of these results to numerical simulations are presented in Fig. 3(a,b).

**FIG. 3.**
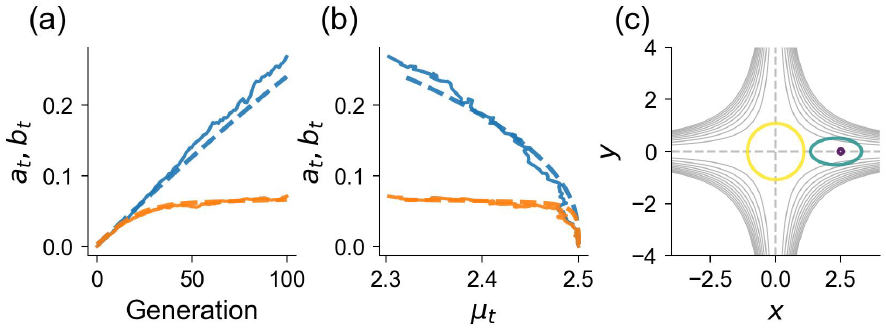
Evolution of population covariance on the degenerate optimal manifold. **(a)** Population variance components *a*_*t*_ (blue) and *b*_*t*_ (orange) as a function of generation. Solid lines show ED simulations; dashed lines show theoretical predictions Eqs. (10–11). **(b)** Same components plotted against mean position *µ*_*t*_ along the degenerate direction, revealing the scaling relationship between moments on the manifold. **(c)** Theoretical covariance ellipses overlaid on the fitness landscape at generations *t* = 0, 100, and 500. The dashed line indicating the optimal manifold (*xy* = 0). In all panels: selection coefficient *β* = 0.1; mutation strength *m*_*x*_ = *m*_*y*_ = 0.05; initial mean ⟨x⟩ _0_ = (2.5, 0)^⊤^; initial covariance Σ_0_ = 0.05 **I**; population size 10,000. Panels (a) and (b) show 100 generations; panel (c) 500 generations of theory.

As time proceeds and the mean drifts towards the origin, the curvature gap between the two directions decreases, and the drift slows down. Finally at long times, the population settles into a steady-state distribution governed by mutation-selection balance; in our approximation this is a Gaussian around the origin with covariance

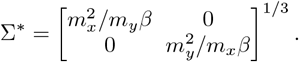

The ratio between the variances reflects only the strength of mutations in the two directions and does not depend on selection, but the individual variances do. Fig. 3(c) depicts the covariance as an ellipse at several points along time, demonstrating the distortion and final steady state.

This analysis illustrates a special case of the more general behavior, where the population variability and curvature interact. The population covariance aligns itself inversely with the Hessian, with larger variance along flatter directions. This provides an anisotropic substrate for mutational search, even if mutations themselves are isotropic, by indirectly encoding the landscape curvature through the population covariance. The result is an effective force that exploits curvature contrasts and pushes the population towards flatter regions. The qualitative generality of this behavior is demonstrated in the Supplementary Material, where alternative selection operators are tested (Sections 1.2), as well as extensions to degenerate manifolds in higher dimensions (Section 5). Effects of a finite population are discussed in Section 2.3.

## E. Generality of Flatness-Seeking Bias

The curvature-driven bias identified above raises a natural question: which aspects of this behavior are specific to evolutionary mutation-selection dynamics, and which arise more generally in stochastic dynamical systems on degenerate landscapes? To clarify this, we compare evolutionary dynamics (ED) with two widely used stochastic frameworks: Langevin dynamics and stochastic gradient descent (SGD).

Langevin dynamics is a canonical physical model of stochastic motion in a potential landscapes with additive noise that is independent of the landscape geometry. Here motion along exactly flat directions is neutral, and no intrinsic directional bias arises. Curvature-dependent drift terms can formally appear in the discretized dynamics, but they are of higher order in the time step and vanish in the continuous-time limit (Supplementary Material, Section 4). Consequently, Langevin dynamics does not exhibit an intrinsic flatness-seeking bias on degenerate fitness manifolds and unbounded diffusion proceeds indefinitely along flat directions. In particular, no normalizable stationary distribution arises over long time in the presence of infinitely extended flat directions, since there is no confinement mechanism. This is in contrast to ED, where the curvature-dependent drift is generated at first order, producing a motion towards flat regions and ultimately a stable, steady-state confined distribution.

Another point for comparison arises from the field of Machine Learning. Recent work has shown that Stochastic Gradient Descent (SGD) exhibits an implicit bias toward flat minima in highly degenerate loss landscapes [9–13]. This phenomenon has raised much interest and is still under intense study, particularly since it was connected to the generalizability and robustness of solutions achieved with this algorithm. Using a physical “shift-model” approximation [13] for an ensemble of SGD optimizers, we find that flatness-seeking behavior indeed emerges, but through a mechanism qualitatively different from Evolutionary Dynamics (Supplementary Material, Section 3). In SGD, effective noise injects larger fluctuations along sharp directions; trajectories are then rapidly expelled from these regions by the gradient flow. This effective injected noise decreases in flat regions; since there is no external noise (playing the role of continuous mutations in ED), the distribution shrinks as flatter regions are populated.

Figure 4 compares the steady-state distributions generated by long-time simulations of ED (a), Langevin dynamics (b), and SGD (c) on the same two-dimensional degenerate landscape (gray contour lines). Although all three systems concentrate near the flattest region, their distributions differ qualitatively, reflecting the distinct ways in which stochasticity interacts with landscape geometry. ED balances mutation and selection to form a confined distribution reflecting these forces; Langevin dynamics uniformly fill the degenerate manifold *xy* = 0, bounded only by the region of simulation; and SGD reduces to practically a point in the flattest region. These comparisons show that flatness-seeking is not a universal consequence of stochasticity, but depends sensitively on how noise couples to the landscape. In evolutionary dynamics, this coupling produces a robust, first-order drift that renders degeneracy an active, directional regime rather than a neutral one.

**FIG. 4.**
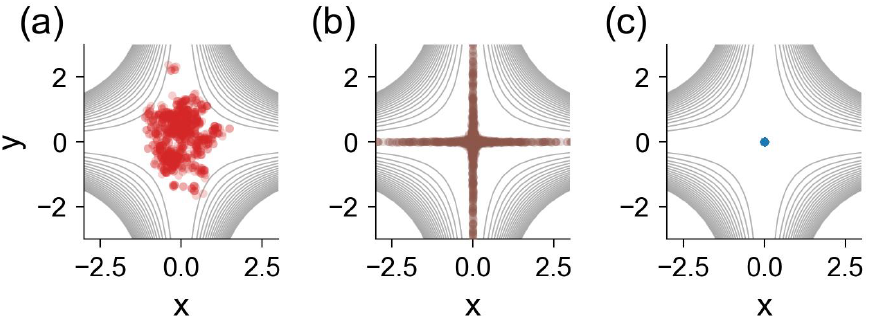
Evolutionary steady-state distributions compared to other stochastic dynamics. **(a)** Evolutionary Dynamics admits a confined steady-state of mutation-selection balance around the flattest point. Gaussian mutation kernel **Σ**_𝕄_ = *σ*^2^**I**, linear selection with intensity *β*. **(b)** Gradient Langevin Dynamics with additive isotropic Gaussian noise fills the flat manifold uniformly with no effective constraining forces. **x**_*t*+1_ = **x**_*t*_ + *η* ∇ *ℱ* (**x**_*t*_) + *ξ*_*t*_. **(c)** Shift model approximating Stochastic Gradient Ascent. **x**_*t*+1_ = **x**_*t*_ + *η*∇ℱ (**x**_*t*_ + *ξ*_*t*_). In **(b)** and **(c)**, *ξ*_*t*_ ∼ 𝒩 (**0**, *σ*^2^**I**). Each panel shows the final population distribution from simulations (scatter) after 10,000 iterations starting from a population uniformly distributed in [−3, 3]^2^. Parameters: population size 1,000; noise/mutation strength *σ* = 0.05; selection intensity/learning rate *β* = *η* = 0.1.

## I. DISCUSSION

In summary, we have explored evolutionary dynamics on a minimal model landscape that captures two essential nontrivial features, continuous degeneracy and heterogeneous curvature. We have shown that, in addition to climbing fitness gradients, evolution is generally subject to an effective force that implicitly induces a directed bias towards flatter regions of phenotype space. On degenerate manifolds, where gradients vanish, this becomes the primary force acting on the population. It arises from curvature differences across trait directions and operates by morphing the standing population variation, extending it preferentially towards flatter regions and thereby allowing mutations to explore these regions more effectively.

Evolution on flat fitness landscapes was previously described as neutral diffusion driven by mutations [16]. Our results show that this picture depends crucially on geometric uniformity: when curvature varies along a degenerate fitness manifold, the nature of dynamics changes dramatically and degeneracy no longer implies neutrality.

Previous work has already demonstrated that degeneracy does not necessarily imply neutrality in discrete settings. In networks of equally fit genotypes, populations concentrate in regions of higher connectivity [17, 18]. Connectivity in a discrete network can be viewed as the analogue of curvature in a continuous landscape: highly connected nodes have more viable neighbors, just as states in flat regions have more locally accessible high-fitness directions.

Holey landscape theory [19] emphasized that in high-dimensional spaces, well-adapted genotypes form extended connected sets that enable evolutionary motion without crossing deep fitness valleys. The smooth landscape studied here was originally proposed in the context of machine learning theory, as a simplified model for high-dimensional degenerate landscapes [11], and enables to develop intuition about the effects of curvature. One can view Holey landscapes as an extreme case where curvature is either zero (between neighboring fit states) or infinity (between fit and unfit), while our model allows the entire spectrum in between. Given the broad distribution of curvatures observed empirically in biological models, “sloppy systems” [20], this is expected to be a useful model that could describe more realistic features of biological landscapes.

Our analysis relies on a mapping over generations that tracks the average and covariance of traits, neglecting higher order statistics. The mapping reformulates classical results in evolutionary theory [21–23], and is related to the family of algorithms of Evolutionary Strategy [24, 25]. We find that already in this approximation it can shed light on the interplay between variability in the population and curvature in the landscape, revealing the implicit bias and drift towards flatter regions, as well as the underlying mechanism.

More broadly, degeneracy is a prevalent feature of biological systems [26, 27], that can in principle supports drift among high-dimensional states of equal functionality. For example in neural systems, representational mappings of environmental cues are known to drift over time [28–30]. Recent work has shown that such drift can arise from implicit biases induced by learning on a degenerate landscape [31]. In growing bacteria, degenerate manifolds can underlie persistent phenotypic variation while maintaining growth-and-division functional homeostasis [32]. These examples suggest that drifts on degenerate high-dimensional manifolds while preserving lower-dimensional functionality, may be a general organizing principle across biological scales.

Taken together, our results have profound implications for how evolution and evolved systems are interpreted. Living systems that we observe today, may have spent substantial evolutionary time drifting in degenerate optimal (or near-optimal) phenotype spaces. Observed patterns of variation therefore do not reflect only adaptation and selection, but may also encode geometric features of the underlying landscape, such as curvature heterogeneity. For example, recent empirical work has shown that genetic variation in quantitative traits across taxa is highly anisotropic, with a large ratio between variance in the two leading dimensions [33]. These patterns were shown to be more consistent with holey landscapes than with other common landscapes. Our results provide a complementary geometric and dynamical interpretation of such observations as potentially reflecting gaps in curvature of directions in trait space. Further study of natural variability in connection to our model is an important direction for future work.

Additional implications concern phenotypic variability, which is often attributed to physical or biochemical noise. In contrast, for a system under drift in degenerate manifolds, variability may reflect structured exploration of high-dimensional phenotype space and encode aspects of evolutionary history. As measurements of phenotypes become increasingly high dimensional, analyzing variation in relation to landscape geometry may become both feasible and necessary. In particular, coexistence of variable phenotypes in a constant environment is not contradictory to optimality, and can encode history and landscape structure.

Finally, biological robustness - a hallmark property of living systems - need not necessarily be implemented by dedicated control mechanisms, as is often assumed. Rather, it can emerge spontaneously through evolution by residing in flat regions of phenotypic landscapes [32, 34]. Mutational robustness has been extensively studied from this perspective [17, 18]; phenotypic robustness less so. Extending our study to higher dimensional systems and investigating how robustness emerges and persists represent important directions for future research.

## Acknowledgments

This work was partially supported by the US-Binational Science Foundation Grant 2024215. We are grateful to Omri Barak, Shimon Marom, Ron Meir and Stefano Recanatesi for for helpful discussions and critical reading of the manuscript.

