## Supplementary Material, text and figures for "Evolution on degenerate fitness landscapes is not neutral: curvature drives directional bias"

Razi Fachareldeen and Naama Brenner, Technion

#### Supplementary Material

**4 Comparison to Langevin dynamics** **12**

**5 Simulations in a High-Dimensional Landscape** **13**

**6 Simulation Details** **14**

#### Notation

Throughout this supplement, we use the following notation: the population distribution at generation $t$ is denoted $p(\mathbf{x}, t)$, with mean $\boldsymbol{\theta}_t = \langle \mathbf{x} \rangle_t$ and covariance $\boldsymbol{\Sigma}_t$. The fitness function is $\mathcal{F}(\mathbf{x})$, the Hessian is $H(\mathbf{x}) = \nabla^2 \mathcal{F}(\mathbf{x})$, and the mutation covariance is $\boldsymbol{\Sigma}_{\mathbb{M}}$. Angle brackets $\langle \cdot \rangle_t$ denote expectations with respect to $p(\mathbf{x}, t)$.

#### Stein identities

For a Gaussian random variable $\mathbf{x} \sim \mathcal{N}(\boldsymbol{\theta}, \boldsymbol{\Sigma})$, and $\mathcal{F} : \mathbb{R}^d \to \mathbb{R}$ smooth with integrable derivatives, we use the Stein identities:

$$\langle (\mathbf{x} - \boldsymbol{\theta})\, \mathcal{F}(\mathbf{x}) \rangle = \boldsymbol{\Sigma}\, \langle \nabla \mathcal{F}(\mathbf{x}) \rangle, \tag{1}$$

$$\langle (\mathbf{x} - \boldsymbol{\theta})(\mathbf{x} - \boldsymbol{\theta})^\top \mathcal{F}(\mathbf{x}) \rangle = \boldsymbol{\Sigma}\, \langle \mathcal{F}(X) \rangle + \boldsymbol{\Sigma}\, \langle \nabla^2 \mathcal{F}(X) \rangle \boldsymbol{\Sigma}. \tag{2}$$

### 1 One-generation mapping for average and covariance

In this section we derive the one-generation mapping for the average and covariance of the population under evolutionary dynamics. Mutations are assumed to be additive to the population covariance, and different selection operators are addressed.

#### 1.1 Linear Selection

Under the mapping, Eq. (3) of the main text, selection is applied in an additive manner:

$$\begin{aligned} p(\mathbf{x}, t+1) &= \mathbb{M}\,\mathbb{S}\,p(\mathbf{x}, t) \\ &= \left[1+\beta(\mathcal{F}(\mathbf{x})-\langle\mathcal{F}\rangle_t)\right] p(\mathbf{x}, t) \, * \, \mathcal{N}(0, \mathbf{\Sigma}_{\mathbb{M}}). \end{aligned} \tag{3}$$

We work in the weak-selection regime in which the linearized selection factor remains nonnegative. Assuming bounded fitness $\mathcal{F}_{\min} \leq \mathcal{F}(x) \leq \mathcal{F}_{\max}$ and writing $\Delta\mathcal{F} = \mathcal{F}_{\max} - \mathcal{F}_{\min}$, a sufficient condition for $\mathbb{S}\,p$ to remain a valid (nonnegative) density for all $t$ is

$$0 \leq \beta < 1/\Delta\mathcal{F}.$$

Under this condition $\mathbb{S}$ preserves normalization since $\int(\mathcal{F} - \langle\mathcal{F}\rangle)p\,dx = 0$.

**Average mapping - smoothing as an implicit bias**

To derive the one-generation update rule for the population's average, we note that applying the mutation operator - a convolution with a distribution of zero mean - does not affect the average; therefore,

$$\langle\mathbf{x}\rangle_{t+1} = \langle\mathbf{x}\rangle_t + \beta\langle(\mathbf{x} - \langle\mathbf{x}\rangle_t)\mathcal{F}(\mathbf{x})\rangle_t,$$

yielding a form of Price equation for the average [1]. Approximating the population distribution by a Gaussian and applying the Stein identity (1), we obtain the one-generation mapping for the population average given in the main text:

$$\boxed{\langle\mathbf{x}\rangle_{t+1} = \langle\mathbf{x}\rangle_t + \beta\,\mathbf{\Sigma}_t\nabla\langle\mathcal{F}(\mathbf{x})\rangle_t.} \tag{4}$$

Moving along the gradient of an averaged fitness landscape $\langle\mathcal{F}(\mathbf{x})\rangle$ rather than the raw fitness $\mathcal{F}(\mathbf{x})$ can induce an implicit bias among states of equal fitness. To demonstrate this, let the random variable over which we average be Gaussian with a general covariance $\mathbf{\Sigma}$ and write $\xi \sim \mathcal{N}(0, \mathbf{\Sigma})$. For small noise, a second-order Taylor expansion gives

$$\mathcal{F}(\mathbf{x} + \xi) \approx \mathcal{F}(\mathbf{x}) + \nabla\mathcal{F}(\mathbf{x})^\top\xi + \tfrac{1}{2}\,\xi^\top H(\mathbf{x})\,\xi, \tag{5}$$

where $\mathbf{H}(\mathbf{x}) = \nabla^2\mathcal{F}(\mathbf{x})$ is the Hessian. Averaging with respect to $\xi$ performs an effective smoothing; the linear term vanishes since $\langle\xi\rangle = 0$, hence

$$\begin{aligned} \langle\mathcal{F}(\mathbf{x} + \xi)\rangle &\approx \mathcal{F}(\mathbf{x}) + \tfrac{1}{2}\,\langle\xi^\top H(\mathbf{x})\,\xi\,\rangle \\ &= \mathcal{F}(\mathbf{x}) + \tfrac{1}{2}\,\mathrm{Tr}(\mathbf{\Sigma}\,H(\mathbf{x}))\,, \end{aligned}$$

where we used the identity (valid for a general noise $\xi$ with covariance $\mathbf{\Sigma}$)

$$\langle\xi^\top\mathbf{A}\xi\rangle = \mathrm{Tr}(\mathbf{\Sigma}\mathbf{A})\,.$$

Applying this to the one-generation update rule for the average, the averaging is performed over the population distribution and we find

$$\langle\mathbf{x}\rangle_{t+1} \;\approx\; \langle\mathbf{x}\rangle_t + \beta\,\mathbf{\Sigma}_t\nabla\Big[\mathcal{F}(\mathbf{x}) + \tfrac{1}{2}\mathrm{Tr}(\mathbf{\Sigma}_t\,H(\mathbf{x}))\Big] \tag{6}$$

To leading order in the population variability. This shows that averaging over the population distribution, effectively a smoothing operation, adds a curvature-dependent bias to the function that is being optimized (appears under the gradient). If the population covariance is aligned along the Hessian eigenbasis, $\mathbf{\Sigma}_t = \mathrm{diag}(\sigma_1^2, \dots, \sigma_d^2)$, then the trace takes the form

$$\mathrm{Tr}(\mathbf{\Sigma}_t\, H(\mathbf{x})) = \sum_{i=1}^{d} \sigma_i^2\, \partial_{ii}\mathcal{F}(\mathbf{x}),$$

i.e., a variance-weighted Laplacian. Near a maximum, $H(\mathbf{x})$ is negative semi-definite, and therefore $\mathrm{Tr}(\mathbf{\Sigma}_t H(\mathbf{x})) \le 0$ (for $\mathbf{\Sigma}_t \succeq 0$), and the implicit bias favors flatter maxima (smaller weighted curvature).

**Covariance matrix update.**

Let $p(\mathbf{x}, t) = \mathcal{N}(\mathbf{x}; \boldsymbol{\theta}_t, \mathbf{\Sigma}_t)$, a Gaussian distribution with mean $\langle \mathbf{x} \rangle_t = \boldsymbol{\theta}_t$ and covariance $\mathbf{\Sigma}_t$. The distribution after selection is

$$\tilde{p}(\mathbf{x}, t+1) = \Big(1 + \beta\big(\mathcal{F}(\mathbf{x}) - \langle \mathcal{F} \rangle_t\big)\Big) p(\mathbf{x}, t)$$

where as usual $\langle \cdot \rangle_t$ denotes expectations with respect to $p(\mathbf{x}, t)$. For any integrable function $g(\mathbf{x})$, its average over $\tilde{p}$ is given by

$$\langle g \rangle_{\tilde{p}} = \langle g \rangle_t + \beta\Big(\langle g\, \mathcal{F} \rangle_t - \langle g \rangle_t \langle \mathcal{F} \rangle_t\Big). \tag{7}$$

Setting $g(\mathbf{x}) = \mathbf{x}$, we recover the update of the average, since the mutation operator does not change the average:

$$\tilde{\boldsymbol{\theta}}_t = \boldsymbol{\theta}_t + \beta \left\langle (\mathbf{x} - \boldsymbol{\theta}_t)\, \mathcal{F}(\mathbf{x}) \right\rangle_t. \tag{8}$$

To compute the post-selection covariance, we apply (7) with $g(\mathbf{x}) = \mathbf{x}\mathbf{x}^\top$:

$$\langle \mathbf{x}\mathbf{x}^\top \rangle_{\tilde{p}} = \mathbf{\Sigma}_t + \boldsymbol{\theta}_t\boldsymbol{\theta}_t^\top + \beta\Big(\langle \mathbf{x}\mathbf{x}^\top \mathcal{F} \rangle_t - (\mathbf{\Sigma}_t + \boldsymbol{\theta}_t\boldsymbol{\theta}_t^\top)\langle \mathcal{F} \rangle_t\Big). \tag{9}$$

The key term $\langle \mathbf{x}\mathbf{x}^\top \mathcal{F} \rangle_t$ is evaluated using the Stein identities. Writing $\mathbf{u} = \mathbf{x} - \boldsymbol{\theta}_t$ and applying (1)–(2) gives

$$\Big\langle \mathbf{u}\, \mathcal{F} \Big\rangle_t = \mathbf{\Sigma}_t \big\langle \nabla \mathcal{F} \big\rangle_t, \tag{10}$$

$$\Big\langle \mathbf{u}\mathbf{u}^\top\, \mathcal{F} \Big\rangle_t = \mathbf{\Sigma}_t \langle \mathcal{F} \rangle_t + \mathbf{\Sigma}_t \big\langle \nabla^2 \mathcal{F} \big\rangle_t \mathbf{\Sigma}_t. \tag{11}$$

Expanding $\mathbf{x}\mathbf{x}^\top = (\boldsymbol{\theta}_t + \mathbf{u})(\boldsymbol{\theta}_t + \mathbf{u})^\top$ and substituting yields

$$\langle \mathbf{x}\mathbf{x}^\top \mathcal{F} \rangle_t = (\mathbf{\Sigma}_t + \boldsymbol{\theta}_t\boldsymbol{\theta}_t^\top)\langle \mathcal{F} \rangle_t + \Delta_t\boldsymbol{\theta}_t^\top + \boldsymbol{\theta}_t\Delta_t^\top + \mathbf{\Sigma}_t \langle H \rangle_t \mathbf{\Sigma}_t, \tag{12}$$

$\Delta_t := \beta\, \mathbf{\Sigma}_t \nabla_{\boldsymbol{\theta}} \langle \mathcal{F} \rangle_t$ is the mean shift from (8). Substituting (12) into (9), the terms $(\mathbf{\Sigma}_t + \boldsymbol{\theta}_t\boldsymbol{\theta}_t^\top)\langle \mathcal{F} \rangle_t$ cancel, giving

$$\langle \mathbf{x}\mathbf{x}^\top \rangle_{\tilde{p}} = \mathbf{\Sigma}_t + \boldsymbol{\theta}_t\boldsymbol{\theta}_t^\top + \beta\, \mathbf{\Sigma}_t \langle \mathbf{H} \rangle_t \mathbf{\Sigma}_t + \Delta_t\boldsymbol{\theta}_t^\top + \boldsymbol{\theta}_t\Delta_t^\top.$$

Since $\boldsymbol{\theta}_{t+1} = \boldsymbol{\theta}_t + \Delta_t$, we have $\boldsymbol{\theta}_{t+1}\boldsymbol{\theta}_{t+1}^\top = \boldsymbol{\theta}_t\boldsymbol{\theta}_t^\top + \Delta_t\boldsymbol{\theta}_t^\top + \boldsymbol{\theta}_t\Delta_t^\top + \Delta_t\Delta_t^\top$, and therefore the post-selection covariance is

$$\tilde{\mathbf{\Sigma}}_t = \mathbf{\Sigma}_t + \beta\, \mathbf{\Sigma}_t \langle \mathbf{H} \rangle_t \mathbf{\Sigma}_t - \Delta_t\Delta_t^\top. \tag{13}$$

Substituting $\Delta_t = \beta\, \mathbf{\Sigma}_t \nabla_{\boldsymbol{\theta}} \langle \mathcal{F} \rangle_t$,

$$\tilde{\mathbf{\Sigma}}_t = \mathbf{\Sigma}_t + \beta\, \mathbf{\Sigma}_t \Big(\langle \mathbf{H} \rangle_t - \beta\, \nabla_{\boldsymbol{\theta}} \langle \mathcal{F} \rangle_t\, \nabla_{\boldsymbol{\theta}} \langle \mathcal{F} \rangle_t^\top\Big) \mathbf{\Sigma}_t. \tag{14}$$

Finally, adding the mutation covariance yields the full one-generation covariance mapping:

$$\boxed{\mathbf{\Sigma}_{t+1} = \mathbf{\Sigma}_t + \beta\, \mathbf{\Sigma}_t \Big(\langle \mathbf{H} \rangle_t - \beta\, \nabla_{\boldsymbol{\theta}} \langle \mathcal{F} \rangle_t\, \nabla_{\boldsymbol{\theta}} \langle \mathcal{F} \rangle_t^\top\Big) \mathbf{\Sigma}_t + \mathbf{\Sigma}_{\mathbb{M}}.} \tag{15}$$

To leading order in $\beta$, this is Eq. 9 of the main text.

###### Steady state covariance matrix

in leading order, the covariance update Eq. (15) can be written as:

$$\mathbf{\Sigma}_{t+1} \approx \mathbf{\Sigma}_t + \beta\, \mathbf{\Sigma}_t \langle \mathbf{H} \rangle_t \mathbf{\Sigma}_t + \mathbf{\Sigma}_{\mathbb{M}}. \tag{16}$$

At steady state we have in this approximation

$$\beta\, \mathbf{\Sigma}^* \langle \mathbf{H} \rangle \mathbf{\Sigma}^* + \mathbf{\Sigma}_{\mathbb{M}} = 0.$$

For an isotropic mutation kernel $\mathbf{\Sigma}_{\mathbb{M}} = \sigma_{\mathbb{M}}^2 \mathbf{I}$ in the eigenbasis of the Hessian, we have

$$\mathbf{\Sigma}^* \langle \mathbf{H} \rangle \mathbf{\Sigma}^* = -\frac{\sigma_M^2}{\beta} \mathbf{I}$$

and multiplying by $(\mathbf{\Sigma}^*)^{-1}$ from the right and left

$$\langle \mathbf{H} \rangle = -\frac{\sigma_{\mathbb{M}}^2}{\beta} (\mathbf{\Sigma}^*)^{-2}, \qquad \mathbf{\Sigma}^* = -\frac{\sigma_{\mathbb{M}}^2}{\beta} \langle \mathbf{H} \rangle^{-1/2}. \tag{17}$$

Since the matrices commute,

$$\mathbf{\Sigma}^* \langle \mathbf{H} \rangle = \langle \mathbf{H} \rangle \mathbf{\Sigma}^* = -\frac{\sigma_{\mathbb{M}}^2}{\beta} (\mathbf{\Sigma}^*)^{-1},$$

they are simultaneously diagonalizable by the same orthogonal matrix $\mathbf{U}$, implying that they have the same eigenvectors:

$$\mathbf{\Sigma} = \mathbf{U}\mathbf{D}_1\mathbf{U}^\mathsf{T}, \qquad \langle \mathbf{H} \rangle = \mathbf{U}\mathbf{D}_2\mathbf{U}^\mathsf{T}$$

and hence we can rewrite (17) in terms of the diagonal matrices,

$$\mathbf{D_1} = \left( -\frac{\sigma_{\mathbb{M}}^2}{\beta} \mathbf{D_2} \right)^{-1/2}. \tag{18}$$

This shows that each steady-state covariance eigenvalue decreases as the square-root of the corresponding (negative) averaged Hessian eigenvalue.

#### 1.2 Alternative Selection Operators

We now generalize the one-generation mapping to selection operators of the form

$$\mathbb{S}\, p(\mathbf{x}, t) = \frac{W(\mathbf{x})}{\langle W(\mathbf{x}) \rangle_t}\, p(\mathbf{x}, t), \tag{19}$$

where $W(\mathbf{x}) \geq 0$ is a weighting function. The post-selection average of any observable $g(\mathbf{x})$ is

$$\langle g(\mathbf{x}) \rangle_{\tilde{p}} = \frac{\langle g(\mathbf{x})\, W(\mathbf{x}) \rangle_t}{\langle W(\mathbf{x}) \rangle_t}. \tag{20}$$

We derive the mean and covariance updates in full generality; specific selection operators then reduce to evaluating $\nabla \ln \langle W \rangle_t$ and $\nabla^2 \ln \langle W \rangle_t$ for the chosen $W$.

###### Mean Update

Since the mutation operator is a zero-mean convolution, the change in the population average is driven entirely by selection. Setting $g(\mathbf{x}) = \mathbf{x}$ in (20) and using the covariance identity $\langle \mathbf{x} W \rangle_t = \langle \mathbf{x} \rangle_t \langle W \rangle_t + \langle (\mathbf{x} - \langle \mathbf{x} \rangle_t) W \rangle_t$:

$$\langle \mathbf{x} \rangle_{t+1} = \langle \mathbf{x} \rangle_t + \frac{1}{\langle W \rangle_t} \langle (\mathbf{x} - \langle \mathbf{x} \rangle_t) W(\mathbf{x}) \rangle_t. \tag{21}$$

Under the Gaussian approximation $\mathcal{N}(\boldsymbol{\theta}_t, \boldsymbol{\Sigma}_t)$, with the application of Stein's identity (1),

$$\langle \mathbf{x} \rangle_{t+1} = \langle \mathbf{x} \rangle_t + \boldsymbol{\Sigma}_t \frac{\langle \nabla W(\mathbf{x}) \rangle_t}{\langle W(\mathbf{x}) \rangle_t}. \tag{22}$$

We recognize the ratio as the gradient of the log-expectation:

$$\boxed{\langle \mathbf{x} \rangle_{t+1} = \langle \mathbf{x} \rangle_t + \boldsymbol{\Sigma}_t \nabla \ln \langle W(\mathbf{x}) \rangle_t.} \tag{23}$$

**Covariance Update**

Setting $g(\mathbf{x}) = \mathbf{x}\mathbf{x}^\top$ in (20) and writing $\mathbf{u} = \mathbf{x} - \boldsymbol{\theta}_t$, Stein's identity gives

$$\langle \mathbf{u}\, W \rangle_t = \boldsymbol{\Sigma}_t \langle \nabla W \rangle_t, \tag{24}$$

$$\langle \mathbf{u}\mathbf{u}^\top W \rangle_t = \boldsymbol{\Sigma}_t \langle W \rangle_t + \boldsymbol{\Sigma}_t \langle \nabla^2 W \rangle_t \, \boldsymbol{\Sigma}_t. \tag{25}$$

Expanding $\mathbf{x}\mathbf{x}^\top = (\boldsymbol{\theta}_t + \mathbf{u})(\boldsymbol{\theta}_t + \mathbf{u})^\top$, substituting these moments, and subtracting the outer product $\tilde{\boldsymbol{\theta}}_t \tilde{\boldsymbol{\theta}}_t^\top$ of the post-selection mean $\tilde{\boldsymbol{\theta}}_t = \boldsymbol{\theta}_t + \Delta_t$ (where $\Delta_t = \boldsymbol{\Sigma}_t \nabla \ln \langle W \rangle_t$), the cross-terms cancel and we obtain the post-selection covariance

$$\tilde{\boldsymbol{\Sigma}}_t = \boldsymbol{\Sigma}_t + \boldsymbol{\Sigma}_t \left( \frac{\langle \nabla^2 W \rangle_t}{\langle W \rangle_t} - \frac{\langle \nabla W \rangle_t \langle \nabla W \rangle_t^\top}{\langle W \rangle_t^2} \right) \boldsymbol{\Sigma}_t. \tag{26}$$

The parenthesized term is precisely $\nabla^2 \ln \langle W \rangle_t$. Adding the mutation covariance $\boldsymbol{\Sigma}_{\mathbb{M}}$:

$$\boxed{\boldsymbol{\Sigma}_{t+1} = \boldsymbol{\Sigma}_t + \boldsymbol{\Sigma}_t \left( \nabla^2 \ln \langle W(\mathbf{x}) \rangle_t \right) \boldsymbol{\Sigma}_t + \boldsymbol{\Sigma}_{\mathbb{M}}.} \tag{27}$$

Equations (23) and (27) show that the entire effect of selection on the Gaussian parameters is encoded in the *log-expected weight* $\ln \langle W \rangle_t$ and its first two derivatives with respect to the mean $\boldsymbol{\theta}_t$. We now evaluate these for two natural choices of $W$.

**Application: Multiplicative Selection**

For $W(\mathbf{x}) = \mathcal{F}(\mathbf{x})$, equations (23)–(27) apply directly with

$$\nabla^2 \ln \langle \mathcal{F} \rangle_t = \frac{\langle H(\mathbf{x}) \rangle_t}{\langle \mathcal{F} \rangle_t} - \frac{\langle \nabla \mathcal{F} \rangle_t \langle \nabla \mathcal{F} \rangle_t^\top}{\langle \mathcal{F} \rangle_t^2}, \tag{28}$$

where $H(\mathbf{x})$ is the Hessian of the fitness function.

**Application: Exponential (Boltzmann) Selection**

For $W(\mathbf{x}) = e^{\mathcal{F}(\mathbf{x})/T}$, we define the *free energy* $\mathcal{E}_t = \ln \langle e^{\mathcal{F}/T} \rangle_t$. The mean update (23) becomes a natural gradient ascent on $\mathcal{E}_t$:

$$\boxed{\langle \mathbf{x} \rangle_{t+1} = \langle \mathbf{x} \rangle_t + \boldsymbol{\Sigma}_t \, \nabla \mathcal{E}_t.} \tag{29}$$

For the covariance update, we need $\nabla^2 \mathcal{E}_t$. Using the chain rule on $W = e^{F/T}$:

$$\nabla W = \tfrac{1}{T} \, W \, \nabla \mathcal{F}, \qquad \nabla^2 W = \tfrac{1}{T} \, W \, \nabla^2 \mathcal{F} + \tfrac{1}{T^2} \, W \, \nabla \mathcal{F} \, \nabla \mathcal{F}^\top. \tag{30}$$

Substituting into the general identity $\nabla^2 \ln \langle W \rangle = \frac{\langle \nabla^2 W \rangle}{\langle W \rangle} - \frac{\langle \nabla W \rangle \langle \nabla W \rangle^\top}{\langle W \rangle^2}$ yields

$$\nabla^2 \mathcal{E}_t = \frac{1}{T} \, \langle H(\mathbf{x}) \rangle_t + \frac{1}{T^2} \, \mathrm{Cov}\big(\nabla \mathcal{F}(\mathbf{x})\big)_t, \tag{31}$$

where the covariance of fitness gradients (Fisher information) arises from the $\nabla F \, \nabla F^\top$ terms. The covariance update (27) therefore reads

$$\boxed{\boldsymbol{\Sigma}_{t+1} = \boldsymbol{\Sigma}_t + \boldsymbol{\Sigma}_t \left[ \frac{1}{T} \, \langle H(\mathbf{x}) \rangle_t + \frac{1}{T^2} \, \mathrm{Cov}\big(\nabla \mathcal{F}(\mathbf{x})\big)_t \right] \boldsymbol{\Sigma}_t + \boldsymbol{\Sigma}_{\mathbb{M}}.} \tag{32}$$

### 2 Two-Dimensional Landscape

#### 2.1 Explicit formulae for the one-generation mapping

Here, we focus on the two dimensional model, where $\mathbf{x} = (x, y)$, and the fitness landscape is

$$\mathcal{F}(x, y) = \mathcal{F}^* - \frac{1}{2}x^2y^2, \tag{33}$$

with gradient and Hessian

$$\nabla\mathcal{F} = -\begin{bmatrix} x\,y^2 \\ y\,x^2 \end{bmatrix}, \qquad H = \nabla^2\mathcal{F} = -\begin{bmatrix} y^2 & 2xy \\ 2xy & x^2 \end{bmatrix}. \tag{34}$$

The population distribution covariance is $\mathbf{\Sigma}_t = \mathrm{diag}(a, b)$, giving the averages over the population

$$\langle\mathcal{F}\rangle = \frac{1}{2}\left[F^* - (x^2 + a)(y^2 + b)\right], \qquad \nabla\langle\mathcal{F}\rangle_t = -\begin{bmatrix} x(y^2 + b) \\ y(x^2 + a) \end{bmatrix} \tag{35}$$

$$\langle H\rangle_t = -\begin{bmatrix} (y^2 + a) & 2xy \\ 2xy & (x^2 + a) \end{bmatrix}. \tag{36}$$

The updates are found as

$$\boldsymbol{\theta}_{t+1} = \boldsymbol{\theta}_t - \beta\,\mathbf{\Sigma}_t \begin{bmatrix} x(y^2 + b) \\ y(x^2 + a) \end{bmatrix}, \tag{37}$$

$$\mathbf{\Sigma}_{t+1} = \mathbf{\Sigma}_t - \beta\,\mathbf{\Sigma}_t \begin{bmatrix} y^2 + b & 2xy \\ 2xy & x^2 + a \end{bmatrix} \mathbf{\Sigma}_t + \mathbf{\Sigma}_{\mathbb{M}} \tag{38}$$

#### 2.2 Dynamics on the degenerate optimal manifold

Here we consider the dynamics on the optimal degenerate manifold under the one-generation mapping. Since the optimal manifold is degenerate with respect to fitness, the dynamics are not driven by fitness gradients but by second-order properties of the manifold. The model landscape disentangles two directions, one flat and one curved with varying curvature. Starting from an initial condition distribution with an average on the $x$-axis and axes-aligned covariance,

$$\boldsymbol{\theta}_t = \begin{bmatrix} \mu_t \\ 0 \end{bmatrix}, \qquad \mathbf{\Sigma}_t = \begin{bmatrix} a_t & 0 \\ 0 & b_t \end{bmatrix},$$

And assuming the mutation matrix is diagonal

$$\mathbf{\Sigma}_{\mathbb{M}} = \begin{bmatrix} m_x & 0 \\ 0 & m_y \end{bmatrix}.$$

The averaged gradient and Hessian on the manifold $y = 0$ takes the form

$$\nabla\langle\mathcal{F}(\boldsymbol{\theta})\rangle = -\begin{bmatrix} \mu_t\,a_t \\ 0 \end{bmatrix}, \qquad \langle H\rangle = -\begin{bmatrix} b_t & 0 \\ 0 & (a_t + \mu_t^2) \end{bmatrix}$$

the update rules are, to leading order in $\beta$,

$$\theta_{t+1} = \begin{bmatrix} \mu_t \\ 0 \end{bmatrix} - \beta \begin{bmatrix} a_t & 0 \\ 0 & b_t \end{bmatrix} \begin{bmatrix} \mu_t b_t \\ 0 \end{bmatrix} = \begin{bmatrix} \mu_t(1 - \beta a_t b_t) \\ 0 \end{bmatrix} \tag{39}$$

$$\mathbf{\Sigma}_{t+1} = \begin{bmatrix} a_t & 0 \\ 0 & b_t \end{bmatrix} - \beta \begin{bmatrix} a_t & 0 \\ 0 & b_t \end{bmatrix} \begin{bmatrix} b_t & 0 \\ 0 & (a_t + \mu_t^2) \end{bmatrix} \begin{bmatrix} a_t & 0 \\ 0 & b_t \end{bmatrix} + \begin{bmatrix} m_x & 0 \\ 0 & m_y \end{bmatrix}. \tag{40}$$

This transformation leaves the covariance matrix diagonal, the average remains on the $x$-axis, and the following update can be written on the three variables defining the distribution $\{\mu_t, a_t, b_t\}$:

$$\mu_{t+1} = \mu_t(1 - \beta a_t b_t) \tag{41}$$
$$a_{t+1} = a_t(1 - \beta a_t b_t) + m_x \tag{42}$$
$$b_{t+1} = b_t(1 - \beta b_t(a_t + \mu_t^2)) + m_y \tag{43}$$

Starting from a region far from the origin with $\mu_t^2 \gg a_t$, the variance in the curved direction $y$ rapidly equilibrates to a quasi-steady-state value

$$0 = -\beta b^2(a + \mu^2)) + m_y \qquad \to \qquad b(a, \mu) = \left[\frac{m_y}{\beta(a + \mu^2)}\right]^{1/2}. \tag{44}$$

with a typical timescale of approximately $\tau_b \sim 1/\beta b_t(\mu_t + a_t)$. Then, the variance in the flat direction follows with a relaxation time $\tau_a \sim 1/\beta a_t b_t$ to its local value. This regime of parameters induces a separation of timescale, where the typical time for the variance in the sharp direction is much faster than the timescale of the flat direction and the average,

$$\frac{\tau_b}{\tau_a} \approx \frac{\beta a_t b_t}{\beta b_t(\mu_t + a_t)} \approx \frac{a_t}{\mu_t} \ll 1.$$

The quasi steady-state values are found as

$$0 = -\beta b a^2 + m_x \qquad \to \qquad a = \left[\frac{m_x}{\beta b(a, \mu)}\right]^{1/2} = \left[\frac{m_x}{\beta}\right]^{1/2}\left[\frac{m_y}{\beta(a + \mu^2)}\right]^{-1/4} \tag{45}$$

Therefore, at large $x$ on the optimal manifold, a strong anisotropy of the population covariance develops:

$$a = \sigma_x^2 \propto |\mu|^{1/2}, \quad b = \sigma_y^2 \propto |\mu|^{-1}, \tag{46}$$

where the second is proportional to the inverse square root of the local curvature in the stiff direction. The first is a much larger spread (the curvature in that direction is zero; $\mu$ is large). At long times, the dynamics reaches a steady-state

$$0 = -\mu^* \beta a^* b^* \tag{47}$$
$$0 = -\beta (a^*)^2 b^* + m_x \tag{48}$$
$$0 = -\beta (b^*)^2 (a_t + \mu_t^2) + m_y \tag{49}$$

assuming that in the presence of mutation $a^* \neq 0, b^* \neq 0$, we get from the first equation $\mu^* = 0$. Solving the other two equations with $\mu^* = 0$,

$$a^* = \sigma_x^2 = \left(\frac{m_x^2}{m_y \beta}\right)^{1/3}, \qquad b^* = \sigma_y^2 = \left(\frac{m_y^2}{m_x \beta}\right)^{1/3}.$$

The ratio $a^*/b^* = m_x/m_y$, independent of selection $\beta$. The actual values of the spread in both directions decreases with selection strength.

#### 2.3 Finite-population effects.

The mean update derived above shows that the population drifts toward the origin at a rate set by $\beta a_t b_t$. In finite populations of size $N$, this deterministic drift competes with sampling noise of order $\sqrt{a_t/N}$ along the flat direction. Using the quasi-steady-state scalings $a_t \propto |\mu_t|^{1/2}$ and $b_t \propto |\mu_t|^{-1}$, the drift rate $\beta a_t b_t \propto |\mu_t|^{-1/2}$ weakens as the population approaches the flattest region, while noise scales as $|\mu_t|^{1/4}/\sqrt{N}$.

To quantify the effects of demographic noise in a finite population, we simulate 100 independent runs for each population size $N$, measuring the signed displacement $D$ toward the origin after $T =$

1000 generations. Figure 1 shows that all population sizes exhibit positive mean drift $\langle D \rangle_N > 0$, with a value increasing as a function of opulation size until a saturation at approximately $N = 100$. The CV of the drift, standard deviation over mean, decreases as expected with $N$. Thus, small populations drift slowly with high variability.

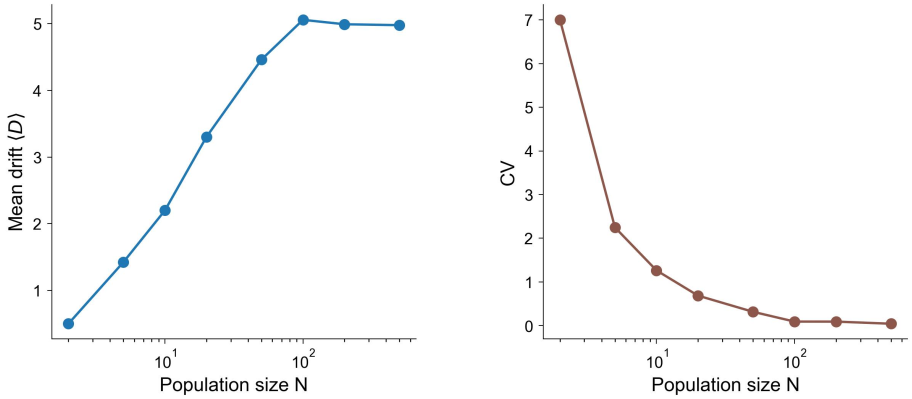

Figure 1: **Finite-population effects on manifold dynamics**. For each population size $N$, 100 simulations were run for $10^4$ generations and the mean and std of the drift $D$ toward the flattest point ($\mu = 0$) was estimated. **(a)** Mean drift $\langle D \rangle$ versus population size $N$. (b) Coefficient of variation; CV $\to 0$ at large $N$ indicates convergence to deterministic dynamics.

#### 2.4 Numerical Validation

We validate the Gaussian-approximation theory derived in previous sections against population-based Monte Carlo simulations for three selection operators: linear (Fig. 2), multiplicative (Fig. 3), and exponential (Fig. 4). In each case, the theoretical results (brown lines) rely on the corresponding mean and covariance update rules: Eqs. (4)–(15) for linear selection, and Eqs. (23)–(27) with $W = F$ or $W = e^{F/T}$ for multiplicative and exponential selection, respectively.

All three figures share the same layout. Panels (a)–(c) show snapshots of the simulated population (red dots) at successive time points, overlaid with the theoretical mean (brown marker) and covariance ellipse (brown outline). The marginal distributions beneath each snapshot compare the empirical $x$-coordinate histogram with the theoretical prediction $\Sigma_{xx}$. Panels (d)–(f) track the dynamics over time: (d) The population mean trajectory on x-y plane; (e) decrease in mean curvature vs time, which continues even after the population has attained maximal fitness; and (f) Fitness vs time.

The Gaussian closure relies on weak selection: the population must remain approximately normal for the second-order moment truncation to hold. As selection pressure increases—a regime more easily accessed by multiplicative and exponential operators than by linear selection—higher-order moments become dynamically relevant, and the approximation degrades. Nevertheless, for the parameter ranges considered here, the Gaussian model robustly captures the essential curvature-driven drift.

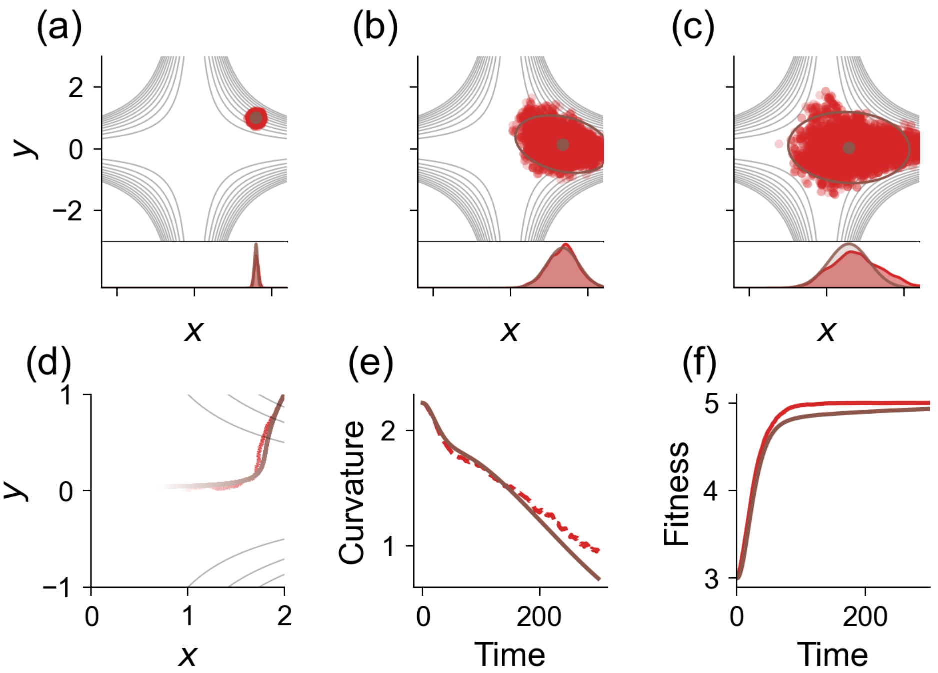

Figure 2: **Linear selection.** Comparison of evolutionary dynamics (ED) simulations with the Gaussian-approximation theory, Eqs. (4)–(15). Parameters: $F^* = 5$, $N = 10{,}000$, initial $\mathbf{\Sigma} = 0.05^2\,\mathbf{I}$, $\eta = \beta = 0.1$. Panels (a)–(c): population snapshots with theoretical mean and covariance ellipse. Panels (d)–(f): mean trajectory in the $x$–$y$ plane, mean curvature, and fitness over time.

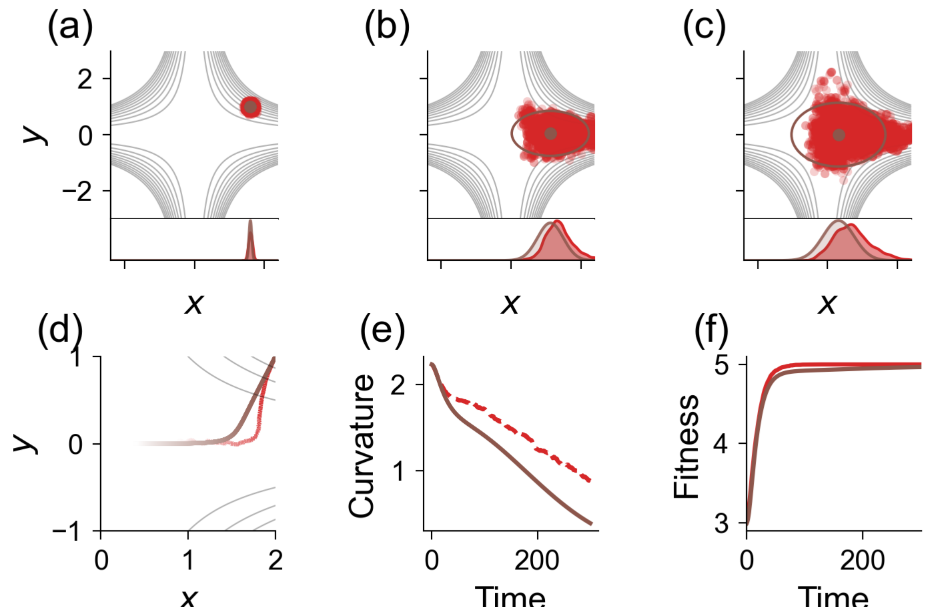

Figure 3: **Multiplicative selection.** Same layout as Fig. 2. The theoretical update implements natural gradient ascent on $\ln\langle F\rangle$ via Eqs. (23)–(27) with $W = F$.

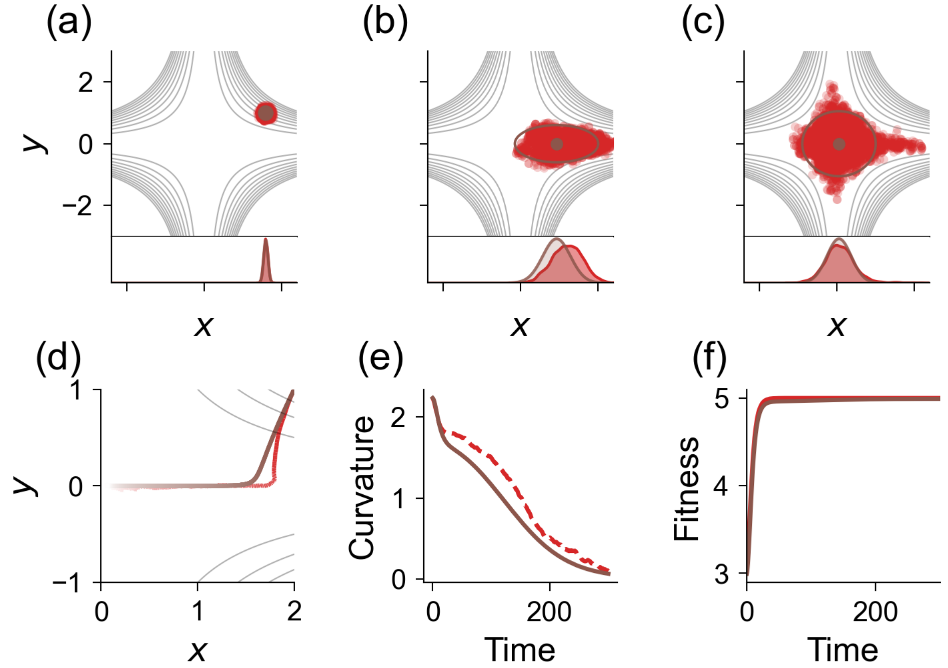

Figure 4: **Exponential selection** ($T = 1$). Same layout as Fig. 2. The theoretical update implements natural gradient ascent on the free energy $\mathcal{E} = \ln\langle e^{F/T}\rangle$ via Eqs. (23)–(27) with $W = e^{F/T}$.

### 3 Comparison to Stochastic Gradient Descent

#### 3.1 One-step mapping for mean and covariance

The shift model [2] describes SGD noise as random perturbations to the parameters $\mathbf{x}$:

$$\nabla^{\mathrm{SGD}}\mathcal{F}(\mathbf{x}) \approx \nabla\mathcal{F}(\mathbf{x}+\boldsymbol{\xi}), \qquad \boldsymbol{\xi} \sim \mathcal{N}(\vec{0}, \boldsymbol{\Sigma}_{sh}) \tag{50}$$

where $\mathbf{x}$ are parameters to be optimized and $\boldsymbol{\Sigma}_{sh}$ denotes the covariance of the shift random variable. For consistency with our evolutionary dynamics model, we consider maximization of a function $\mathcal{F}$ (rather than minimization of a loss function). The one-step mapping is

$$\mathbf{x}_{t+1} = \mathbf{x}_t + \eta\nabla\mathcal{F}(\mathbf{x}_t + \boldsymbol{\xi}_t). \tag{51}$$

where $\eta$ is the step-size, sometimes termed learning rate. To compare with the one-generation formalism developed for ED, we imagine an ensemble of trajectories starting at a Gaussian distribution of parameters $\mathbf{x} \sim \mathcal{N}(\boldsymbol{\theta}_t, \boldsymbol{\Sigma}_t)$. Then, $Z_t = \mathbf{x}_t + \boldsymbol{\xi}_t$ is also Gaussian with the same mean and with a modified covariance $\tilde{\boldsymbol{\Sigma}}_t = \boldsymbol{\Sigma}_t + \boldsymbol{\Sigma}_{sh}$: namely, $Z_t \sim \mathcal{N}(\boldsymbol{\theta}_t, \tilde{\boldsymbol{\Sigma}}_t)$. Using the same notations as our ED one-generation mapping (4), we expand the shift in the gradient to leading order, and derive the one-step mapping:

$$\langle\mathbf{x}\rangle_{t+1} = \langle\mathbf{x}\rangle_t + \eta\,\nabla\langle\mathcal{F}(\mathbf{x})\rangle_t \approx \langle\mathbf{x}\rangle_t + \eta\,\nabla\Big[\mathcal{F}(\mathbf{x}) + \tfrac{1}{2}\mathrm{Tr}\Big(\tilde{\boldsymbol{\Sigma}}_t\,H(\mathbf{x})\Big)\Big]. \tag{52}$$

with the average performed over the distribution of perturbed parameters $Z_t$. We can see that the one-step mapping for SGD is similar in structure to that of ED; the average over the random variable $Z_t$ smooths the landscape and introduces a similar implicit bias. However, in SGD the average proceeds along the *bare* gradient whereas ED follows the *natural* gradient.

For a covariance update, we calculate directly

$$\Sigma_{t+1} = \mathrm{Cov}(\mathbf{x}_\mathrm{t} + \eta\nabla\mathcal{F}(\mathrm{Z}_\mathrm{t})) \tag{53}$$

$$= \Sigma_t + \eta\,\mathrm{Cov}(\mathbf{x}_\mathrm{t}, \nabla\mathcal{F}(\mathrm{Z}_\mathrm{t})) + \eta\,\mathrm{Cov}(\nabla\mathcal{F}(\mathrm{Z}_\mathrm{t}), \mathbf{x}_\mathrm{t}) + \eta^2\,\mathrm{Cov}(\nabla\mathcal{F}(\mathrm{Z}_\mathrm{t})). \tag{54}$$

We first evaluate the covariance

$$\mathrm{Cov}(\mathbf{x}, \mathrm{Z}) = \langle(\mathbf{x}-\boldsymbol{\theta})(\mathrm{Z}-\boldsymbol{\theta})^\top\rangle = \langle(\mathbf{x}-\boldsymbol{\theta})(\mathbf{x}+\boldsymbol{\xi}-\boldsymbol{\theta})^\top\rangle = \Sigma$$

where we have used the fact that $\langle\boldsymbol{\xi}\rangle = 0$. Then, using Stein's conditional identity for covariance,

$$\mathrm{Cov}(\mathbf{x}_\mathrm{t}, \nabla\mathcal{F}(\mathrm{Z}_\mathrm{t})) = \mathrm{Cov}(\mathbf{x}, \mathrm{Z})\langle\mathrm{H}\rangle_\mathrm{t} = \Sigma_\mathrm{t}\langle\mathrm{H}\rangle_\mathrm{t}$$

Finally giving the covariance mapping

$$\Sigma_{t+1} = \Sigma_t + \eta\left[\Sigma_t\langle H\rangle + \langle H\rangle\Sigma_t\right] + \eta^2\Gamma_t, \tag{55}$$

Where we have defined

$$\Gamma_t := \mathrm{Cov}(\nabla\mathcal{F}(\mathrm{Z}_\mathrm{t})) = \langle\nabla\mathcal{F}_\mathrm{t}\nabla\mathcal{F}_\mathrm{t}^\top\rangle - \langle\nabla\mathcal{F}_\mathrm{t}\rangle\langle\nabla\mathcal{F}_\mathrm{t}^\top\rangle.$$

Eq. (55) has a very different structure from the covariance map of ED. We note that, if we keep only the leading order in $\eta$, it admits only the trivial steady-state solution $\Sigma^* = 0$. Keeping both terms in the full update equation, and assuming the matrices are in quasi-steady-state, we find the solution

$$\Sigma^*\langle H\rangle + \langle H\rangle\Sigma^* + \eta\Gamma_t = 0. \tag{56}$$

If they can be diagonalized in the same basis, we find

$$\Sigma^*_{ii} \approx \frac{\eta\Gamma_{ii}}{H_{ii}}. \tag{57}$$

The scaling with the Hessian $\Sigma^* \sim \eta/H$ is very different from the scaling we found for evolutionary dynamics $\Sigma^* \sim 1/H^{1/2}$. Moreover, the steady state covariance is proportional to the step size $\eta$, implying that in the limit of small step size it collapses to a point of the same order as $\eta$. This is in marked contrast to the steady state of ED, which was composed of a balance between mutation and selection.

#### 3.2 Explicit formulae for the optimal manifold

We derive the one-step mapping explicitly for the average and covariance of the parameters on the model landscape $\mathcal{F}(\mathbf{x}) = \mathcal{F}^* - \frac{1}{2}x^2y^2$. Starting from the initial condition $\boldsymbol{\theta}_t$, the SGD equation of motion reads

$$\mathbf{x}_{t+1} = \mathbf{x}_t + \eta\begin{bmatrix}(x_t+\xi_{t,x})\xi_{t,y}^2\\(x_t+\xi_{t,x})^2\xi_{t,y}\end{bmatrix}. \tag{58}$$

Now defining an initial ensemble as above, $\mathbf{x} \sim \mathcal{N}(\boldsymbol{\theta}_t, \Sigma_t)$ and the shift perturbation as

$$\boldsymbol{\theta}_t = \begin{bmatrix}\mu_t\\0\end{bmatrix}, \quad \Sigma_t = \begin{bmatrix}a_t & 0\\0 & b_t\end{bmatrix} \quad \tilde{\Sigma}_t = \begin{bmatrix}\tilde{a}_t & 0\\0 & \tilde{b}_t\end{bmatrix}, \tag{59}$$

We use the update rule and the gradient of this landscape, and average over $Z_t = \mathbf{x}_t + \boldsymbol{\xi}$ to find to leading order in $\eta$

$$\boldsymbol{\theta}_{t+1} = \boldsymbol{\theta}_t + \begin{bmatrix}\mu_t(1-\eta\tilde{b}_t)\\0\end{bmatrix}. \tag{60}$$

Similarly for the covariance update

$$\Sigma_{t+1} = \Sigma_t - 2\eta\begin{bmatrix}\tilde{a}_t\tilde{b}_t & 0\\0 & \tilde{b}_t(\mu_t^2+\tilde{a}_t)\end{bmatrix} + \eta^2\tilde{b}_t\begin{bmatrix}\tilde{b}(2\mu_t^2+3\tilde{a}_t) & 0\\0 & (\mu_t^4+6\mu_t^2\tilde{a}_t+3\tilde{a}_t^2)\end{bmatrix} \tag{61}$$

We see that at large $\mu_t$, where the curvature in the $y$-direction is very high, the variance in that direction is extremely large, $\propto \mu_t^4$, whereas in the flat direction it is only $\propto \mu_t^2$. This is a manifestation of the nature of SGD noise, which is largest in the most curved directions.

### 4 Comparison to Langevin dynamics

The single-particle Langevin stochastic equation of motion can be written as

$$\mathbf{x}_{t+1} = \mathbf{x}_t + \eta\nabla\mathcal{F}(\mathbf{x}_t) + \sigma\,\boldsymbol{\xi}_t. \tag{62}$$

where $\boldsymbol{\xi}_t = (\xi_{t,x}, \xi_{t,y})$ are i.i.d. standard Gaussian (independent across time and coordinates). The noise term can be scaled as a discrete gradient dynamics, with two separate parameters for the time-step and the noise amplitude, or as a physical scaling representing discretizations of a continuous stochastic equation:

$$\sigma \to \begin{cases} \eta\,\sigma, & \text{noisy-gradient dynamics: } \mathbf{x}_{t+1} = \mathbf{x}_t + \eta\big[\nabla\mathcal{F}(\mathbf{x}_t) + \sigma\boldsymbol{\xi}_t\big], \\ \sqrt{2\eta}\,\sigma, & \text{Euler–Maruyama Langevin: } \mathbf{x}_{t+1} = \mathbf{x}_t + \eta\,\nabla\mathcal{F}(\mathbf{x}_t) + \sqrt{2\eta}\,\sigma\,\boldsymbol{\xi}_t. \end{cases} \tag{63}$$

We derive the formulas for a general $\sigma$ and then discuss the scaling at the end. Component-wise on $\mathcal{F} = \mathcal{F}^* - \frac{1}{2}x^2y^2$,

$$x_{t+1} = x_t - \eta\,x_t y_t^2 + \sigma\,\xi_{t,x}, \tag{64}$$

$$y_{t+1} = y_t - \eta\,x_t^2 y_t + \sigma\,\xi t, y. \tag{65}$$

We follow Langevin dynamics on the degenerate optimal manifold by starting from $\mathbf{x}_t = (x_t, 0)^\top$. Since the gradient vanishes on the manifold,

$$x_{t+1} = x_t + \sigma\,\xi_{t,x}, \qquad y_{t+1} = \sigma\,\xi_{t,y}, \tag{66}$$

and on average, as the noise averages to zero, we find that after one time step

$$\langle x\rangle_{t+1} = x_t \tag{67}$$

$$\langle y\rangle_{t+1} = y_t = 0. \tag{68}$$

Namely on average over the noise, the trajectory did not advance from its initial position. At the next time step,

$$\begin{aligned} x_{t+2} &= x_{t+1} - \eta\,x_{t+1}y_{t+1}^2 + \sigma\,\xi_{t+1,x} \\ y_{t+2} &= -\eta\,x_t^2 y_t + \sigma\,\xi_{t,y}. \end{aligned} \tag{69}$$

and on average, since $\langle y\rangle_{t+1}^2 = \sigma^2$

$$\begin{aligned} \langle x\rangle_{t+2} &= x_t - \eta\,x_t\sigma^2 \\ \langle y\rangle_{t+2} &= 0. \end{aligned} \tag{70}$$

Now for the two alternative scaling of the noise amplitudes (63) into (69):

$$\langle x_{t+2} - x_t \mid x_t,\ y_t = 0\rangle = \begin{cases} -\eta^3\sigma^2\,x_t, & \text{(noisy-gradient)}, \\ -2\eta^2\,x_t, & \text{(Langevin)}. \end{cases} \tag{71}$$

In both cases there is a directed drift toward the flattest point $x = 0$ after two discrete steps, namely it is a second-order effect that depends on the variance in the $y$ direction. Depending on the scaling of the noise, the drift has a different scaling : $\mathcal{O}(\eta^3)$ for noise injected inside the gradient step (amplitude $\propto \eta$), and $\mathcal{O}(\eta^2)$ for Langevin noise (amplitude $\propto \sqrt{\eta}$). In both cases, the directed drift is a high order term that vanishes in the continuum limit. The average drift velocity vanishes with the time step

$$\langle v_D\rangle = \frac{\langle x_{t+2} - x_t\rangle}{2\eta} \to 0.$$

### 5 Simulations in a High-Dimensional Landscape

We consider the high-dimensional generalization of the fitness landscape introduced in the main text (Eq. 6):

$$\mathcal{F}(\mathbf{x}, \mathbf{y}) = \mathcal{F}^* - \frac{1}{2}\sum_{i=1}^{n} x_i^2 \lambda_i(\mathbf{y}) \tag{72}$$

where $\mathbf{x} \in \mathbb{R}^n$ represents the coordinates in the non-degenerate (sharp) subspace, and $\mathbf{y} \in \mathbb{R}^{d-n}$ represents the coordinates in the degenerate (flat) subspace. The manifold of optimal solutions corresponds to $\mathbf{x} = 0$. The curvature in each sharp direction $x_i$ is given by the effective eigenvalue $\lambda_i(\mathbf{y})$, which varies with position $\mathbf{y}$ along the degenerate manifold.

To generate diverse landscape instances for the numerical simulations presented in Figs. 5 and 6, we utilized a specific quadratic curvature profile. In the notation of Eq. 72 (where $\mathbf{y}$ denotes the degenerate manifold):

$$\lambda_i(\mathbf{y}) = \lambda_{0,i} + \mathbf{y}^T \boldsymbol{\Lambda}_i \mathbf{y}. \tag{73}$$

Here, $\lambda_{0,i} > 0$ sets the baseline curvature, and the positive semi-definite matrix $\boldsymbol{\Lambda}_i$ determines how curvature increases away from the origin. This construction ensures that the origin $(\mathbf{x} = 0, \mathbf{y} = 0)$ is the unique flattest optimum. The landscape parameters were randomized as follows:

- **Baseline Curvature**: $\lambda_{0,i}$ were drawn uniformly from $[0.01, 1.0]$.
- **Curvature Variation**: The matrices $\boldsymbol{\Lambda}_i \in \mathbb{R}^{(d-n)\times(d-n)}$ were constructed to be symmetric and positive definite. For each $i$, we generated a random orthogonal matrix $\mathbf{Q}$ and a set of random positive eigenvalues, forming $\boldsymbol{\Lambda}_i = \mathbf{Q}\text{diag}(\mu_1, \ldots, \mu_{d-n})\mathbf{Q}^T$. This ensures that the landscape curvature is always positive in the sharp directions and varies smoothly across the flat manifold.

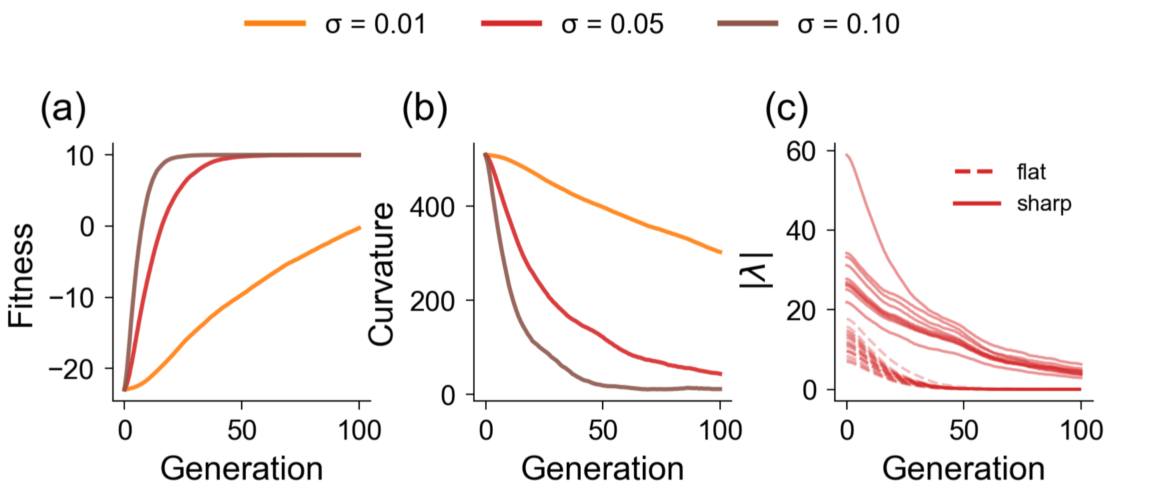

Figure 5: **High-dimensional evolutionary dynamics with varying mutation strength.** Fitness landscape has $N_F = 20$ flat and $N_S = 10$ sharp directions ($D = 30$). The mutation kernel is isotropic Gaussian, $\Sigma_{\mathbb{M}} = \sigma^2\mathbf{I}$, and selection is linear with $\beta = 0.1$. **(a)** Mean fitness vs generation for $\sigma = \{0.01, 0.05, 0.1\}$. **(b)** Mean curvature $|\text{Tr}(H(\langle\mathbf{x}\rangle_t))|$ along the same trajectories. **(c)** Eigenvalue magnitudes $|\lambda_i(t)|$ for the middle mutation strength ($\sigma = 0.05$), plotted for all $D$ directions and grouped by flat (dashed) vs sharp (solid) eigenmodes. Population size $N = 5000$; 100 generations.

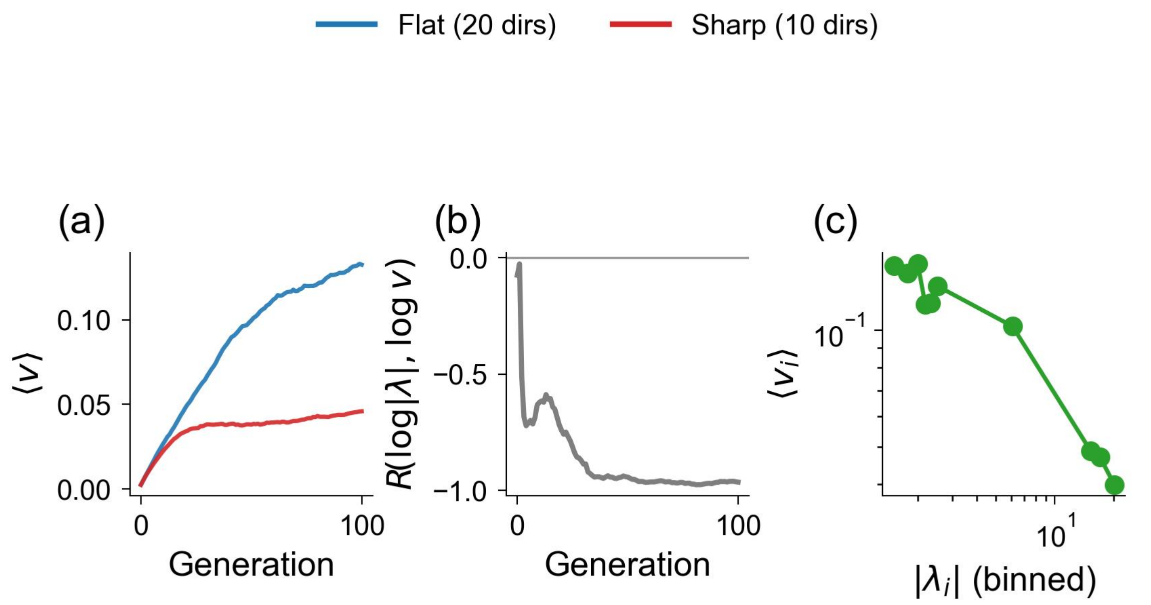

Figure 6: **Variance–curvature structure during high-D evolution.** Dynamics are from the $\sigma = 0.05$ run in the $D = 30$ landscape with $N_F = 20$ flat and $N_S = 10$ sharp directions. **(a)** Mean projected variance onto flat vs sharp subspaces, $\langle v \rangle_{\text{flat}}$ and $\langle v \rangle_{\text{sharp}}$. **(b)** Pearson correlation $R(\log|\lambda|, \log v)$ between Hessian eigenvalue magnitudes and projected variances across directions. **(c)** Final-time binned trend: mean projected variance $\langle v_i \rangle$ vs $|\lambda_i|$ (log–log), with ticks shown as powers of ten. Population size $N = 5000$; 100 generations; selection $\beta = 0.1$.

The high-dimensional simulations confirm the theoretical predictions developed in the 2D setting. Fig. 5 shows that the population drifts toward flatter regions of the manifold (decreasing curvature over time), with larger mutation rates driving faster and more pronounced flatness-seeking dynamics. Fig. 6 directly tests the variance–curvature relationship derived in Section 2. The population covariance aligns with the Hessian eigenbasis and exhibits a strong negative correlation between projected variance and curvature magnitude. Panel (c) confirms the scaling $\sigma_i^2 \propto |\lambda_i|^{-1/2}$ predicted by the steady-state covariance analysis (Eq. 17), demonstrating that the "spectral adaptation" mechanism—where the population spreads more along flat directions and compresses along sharp ones—generalizes robustly to high dimensions.

#### 6 Simulation Details

While the main text presents evolution as operations on continuous distributions, our numerical implementation uses a particle-based approach to simulate the dynamics. Here we detail the algorithm and its correspondence to the theoretical framework.

The population distribution $p(\mathbf{x}, t)$ is approximated by a finite collection of $N$ individuals:

$$p(\mathbf{x}, t) \approx \frac{1}{N} \sum_{i=1}^{N} \delta(\mathbf{x} - \mathbf{x}_i^{(t)}) \tag{74}$$

where $\mathbf{x}_i^{(t)}$ denotes the position of individual $i$ at generation $t$. This discrete representation allows efficient simulation while converging to the continuous dynamics as $N \to \infty$.

Each generation proceeds through the following steps:

**Algorithm 1** Evolutionary Dynamics Simulation

**Input:** Initial mean $\boldsymbol{\mu}_0$, population size $N$, iterations $T$, mutation covariance $\boldsymbol{\Sigma}_{\mathbb{M}}$, selection strength $\beta$
**Initialize:** Sample population $\{\mathbf{x}_i^{(0)}\}_{i=1}^N \sim \mathcal{N}(\boldsymbol{\mu}_0, \boldsymbol{\Sigma}_0)$
**for** $t = 0$ to $T - 1$ **do**
  **Mutation:** For each $i$, sample $\mathbf{x}'_i \sim \mathcal{N}(\mathbf{x}_i^{(t)}, \boldsymbol{\Sigma}_{\mathbb{M}})$
  **Fitness Evaluation:** Compute $f_i = \mathcal{F}(\mathbf{x}'_i)$ for all $i$
  **Selection:**
  - Calculate mean fitness: $\bar{f} = \frac{1}{N}\sum_i f_i$
  - Compute selection probabilities: $p_i = \max(0, 1 + \beta(f_i - \bar{f}))$
  - Normalize: $p_i \leftarrow p_i / \sum_j p_j$
  - Sample with replacement: $\{\mathbf{x}_i^{(t+1)}\}_{i=1}^N \sim \text{Multinomial}(\{\mathbf{x}'_i\}, \{p_i\})$

**end for**

- **Mutation Operator**: The mutation step implements the convolution $\mathbb{M}p(\mathbf{x}, t)$ by adding independent Gaussian noise to each individual:

$$\mathbf{x}'_i = \mathbf{x}_i^{(t)} + \boldsymbol{\epsilon}_i, \quad \boldsymbol{\epsilon}_i \sim \mathcal{N}(0, \boldsymbol{\Sigma}_{\mathbb{M}}) \tag{75}$$

  This is equivalent to sampling from the kernel $\mathcal{N}(\mathbf{x}|\mathbf{x}_i^{(t)}, \boldsymbol{\Sigma}_{\mathbb{M}})$.

- **Selection Operator**: The linear selection operator from Eq. (**??**) is implemented through Resampling (with replacement) with probabilities proportional to the selection weights. The normalization step ensures $\sum_i p_i = 1$, and the $\max(0, \cdot)$ operation handles cases where $\beta(f_i - \bar{f}) < -1$, preventing negative probabilities.
